# Genomic evaluation of a case of *Lacticaseibacillus rhamnosus* endocarditis: comparison with commercial yogurt isolates and examination of global population structure

**DOI:** 10.1101/2022.12.13.520361

**Authors:** Phillip P. Santoiemma, Susan E. Cohn, Samuel W. M. Gatesy, Alan R. Hauser, Saaket Agrawal, Maria Theodorou, Kelly E.R. Bachta, Egon A. Ozer

## Abstract

Probiotics, including yogurt commonly containing bacteria of the *Lactobacillus* group, are widely used to promote microbiome health and to prevent gastrointestinal or other infections, especially in the setting of antibiotic use. Rarely, however, *Lactobacillus* bacteria have caused invasive or severe infections in patients. In this report, we present a patient with fatal *Lacticaseibacillus* bacteremia and endocarditis who frequently consumed yogurt. Whole genome sequencing identified the patient’s bloodstream isolate as *Lacticaseibacillus* (formerly *Lactobacillus*) *rhamnosus*, and sequencing of *Lactobacillus* isolates from selected commercial yogurt preparations revealed one isolate nearly identical to the patient’s isolate, differing by only one nucleotide (adenine to guanine substitution leading to a M1A amino acid change in the start codon of the *nrdR* gene). To place these sequences into broader context of *L. rhamnosus* strains, we compared them to 602 *L. rhamnosus* genome sequences publicly available through the National Center for Biotechnology Information repository. Both isolates belonged to a clade, identified in this report as clade YC, composed of mostly gastrointestinal isolates from healthy individuals, some of which also differed by only a single nucleotide from the patient’s isolate. Members of clade YC had only rarely been previously identified in invasive infections. In summary, the *Lacticaseibacillus* strain causing this patient’s infection may have originated from a commercial dairy product or from the commensal flora of his gastrointestinal tract. This study demonstrates that whole genome sequencing may be insufficient to reliably determine the source of invasive infections caused by *L. rhamnosus*.

**IMPORTANCE:** Active culture and probiotic foods and supplements are often used to promote gut health. Though largely considered safe for human consumption, there are rare reports of severe infections linked to bacterial species used in these products. Using whole genome sequencing, we examined a case of fatal infection with *Lacticaseibacillus rhamnosus* in the patient’s blood. By comparing the bacterial genome sequence from the patient to strains cultured from 4 commercial yogurt samples and 602 publicly available sequences of *L. rhamnosus* from around the world, we found that it was nearly identical to one of the yogurt strains and equally similar to other strains from both probiotics and healthy individuals’ gastrointestinal tracts. These findings show that even with whole genome sequencing it can be difficult to determine the source of an invasive *L. rhamnosus* infection, whether from recent consumption of a probiotic or dairy product versus from the gastrointestinal microbiome.

## INTRODUCTION

In recent years, probiotics have been increasingly used to promote general health and prevent infections such as *Clostridioides difficile* colitis, bacterial vaginosis, and ventilator-associated pneumonia [1]. Particular attention has been given to species in the *Lactobacillus* group, which are found in fermented dairy products. For this reason, yogurt has been promoted as a beneficial source of *Lactobacillus* probiotics [2]. Although these bacteria are generally considered non-pathogenic and safe, more than 200 cases of severe *Lactobacillus* infections have been reported, including bloodstream infections and infective endocarditis [3–6]. Of particular interest is whether ingested *Lactobacillus*-containing foods or probiotic products can escape the gastrointestinal (GI) tract to cause invasive infections. This is especially pertinent to patients with underlying disease conditions or who undergo diagnostic and surgical procedures that compromise the integrity of the GI tract. Confounding this is the fact that several species of *Lactobacillus* are considered part of the normal flora of the GI and female genital tracts [7]. Therefore, it is possible that commensal rather than ingested bacteria may serve as a source of infection.

Whole genome sequencing (WGS) approaches are increasingly being applied to molecular epidemiological investigations of infections and outbreaks. WGS-based typing of cultured microbes allows for higher resolution and discrimination of isolates than older techniques such as pulsed field gel electrophoresis (PFGE) or random amplification of polymorphic DNA (RAPD), both of which are prone to both over- and underestimation of isolate relatedness and diminished inter-laboratory reproducibility [8, 9]. Examining variation across the entire genome with WGS can detect more discriminatory features than other sequence-based typing methods such as multi-locus sequence typing (MLST). MLST is limited to assessing variability across just seven conserved genes and MLST databases are often not well-populated or publicly available for many rare or emerging pathogens. In contrast, WGS provides high resolution characterization of variability among clinical isolates and can readily be compared to available sequencing data generated at any other institution worldwide. This portability of high-resolution WGS data, combined with relevant metadata such as culture dates, geographic location, or isolation source, can precisely define strain relatedness as well as the local or global context of isolates under investigation.

Here, we present a follow-up study to a previously reported case of fatal *Lactobacillus* endocarditis in a patient who consumed yogurt daily [10]. WGS further identified the isolate from the patient’s bloodstream as *Lacticaseibacillus* (formerly *Lactobacillus*) *rhamnosus*, a common human and animal commensal species as well as one used frequently in commercial products. Comparative WGS revealed that the *L. rhamnosus* bloodstream isolate was nearly identical to *L. rhamnosus* bacteria cultured from one of four samples of commercially available yogurts, potentially linking the infection to yogurt consumption. However, the sequences of the yogurt and bloodstream isolates were also nearly identical to many previously sequenced *L. rhamnosus* isolates obtained from the GI tracts of healthy patients. These results indicate that comparative genomic analysis may be insufficient to discriminate between potential sources of *L. rhamnosus* infections.

## CASE REPORT

In December 2019, an 83-year-old male presented with fever, cough and rigors. His past medical history was significant for transaortic valve replacement for aortic stenosis in 2017, osteoarthritis and lumbar stenosis requiring posterior spinal fusion in 2013, Crohn’s Disease requiring small bowel resection in 2000, hepatitis C infection mediated cirrhosis followed by sustained virologic response after treatment, IgG monoclonal gammopathy treated with chronic intravenous immunoglobulin (IVIG) infusions and prostate cancer in remission. A few weeks prior to presentation, the patient underwent a dental cleaning for which he took prophylactic amoxicillin. On presentation, the patient was diagnosed with vertebral osteomyelitis and *Lactobacillus* bacteremia but had no evidence of endocarditis by transesophageal echocardiogram (TEE). He was treated with intravenous (IV) ampicillin for 6 weeks. Two weeks after completion of this antibiotic course, he presented with a second episode of prolonged *Lactobacillus* bacteremia marked by five consecutive days of positive blood cultures. On repeat TEE, he was found to have a vegetation on his prosthetic aortic valve. He was treated with IV ampicillin and gentamicin and initially improved but one week later, developed a large parietal intracranial hematoma presumably from rupture of a mycotic aneurysm. He subsequently decompensated, was transitioned to comfort care and passed away in March 2020. A more extensive report of this patient’s clinical case has been previously published outlining further details of his clinical course [10].

The patient and his family indicated to the clinical staff that he regularly ate yogurt to promote GI health and his family later identified what they believed to be his favorite brands. Given that the *Lactobacillus* group of bacteria are commonly used as active culture agents in commercial yogurt products, we investigated whether there could be a link between his regular yogurt consumption and his invasive *Lactobacillus* infection. We performed comparative WGS of the patient’s bloodstream isolate and *Lactobacillus* isolates cultured from four brands of commercially available yogurt.

## RESULTS

To identify the species of bacterium causing the patient’s infection, the strain isolated from the patient’s bloodstream during his second hospitalization was whole genome sequenced. Additionally, we purchased single servings of four different yogurt brands (anonymized in this report as A, B, C, and D) from a local branch store of a North American supermarket chain. Samples of each yogurt were cultured under *Lactobacillus-*selective conditions, and six cultured colonies from each brand were chosen at random (designated A-1 through 6, B-1 through 6, etc.) for whole genome sequencing. Two of the isolates selected, A-3 and B-1, sequenced poorly and were excluded from further analyses. Analysis of the 16S rRNA gene sequences from the genome assemblies identified the patient’s bloodstream isolate, as well all isolates from brands A and D and three of the six isolates from brand C as *Lacticaseibacillus rhamnosus*. The other three isolates from brand C and all six isolates from brand B speciated as *Lacticaseibacillus paracasei*. Reference-based read alignment and phylogenetic analysis of the *L. rhamnosus* yogurt isolates revealed a clonal relationship among isolates from brands A and D (Fig. 1). The three *L. rhamnosus* isolates from brand C and the patient’s bloodstream isolate were also found to be clonal with each other. Due to a lack of genetic variability among isolates of the same species within each yogurt sampled, isolates A-1, C-2, and D-1 were selected as representative *L. rhamnosus* isolates from each yogurt brand.

**Figure 1.**
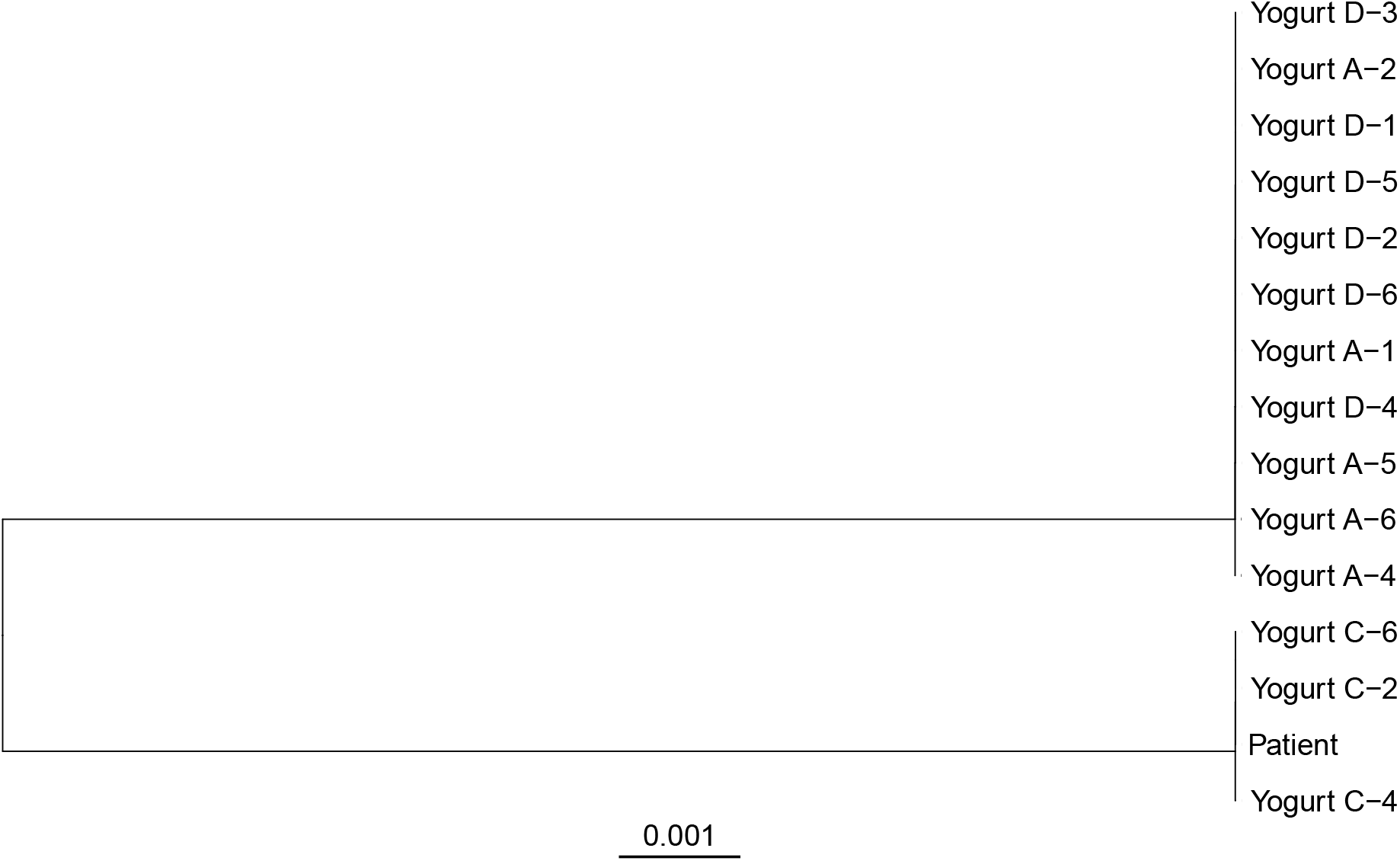
Maximum likelihood whole-genome phylogenetic tree of *Lacticaseibacillus rhamnosus* sequences from a patient with bloodstream infection and sampled yogurt brands. Sequencing reads were aligned to the genome sequence of strain ATCC 11443 and the maximum likelihood phylogenetic tree was determined from the whole genome sequence alignment. For yogurt isolates, letters (A, C, D) indicate separate yogurt brands sampled and numbers (1 – 6) indicate separate colonies cultured from each yogurt brand. Scale bar indicates genetic distance.

To contextualize the patient and yogurt isolates within the larger population of *L. rhamnosus*, we compared these sequences to other publicly available *L. rhamnosus* genome sequences and associated metadata. We queried the assembly and short read archive (SRA) databases of the National Center for Biotechnology Information (NCBI) to identify unique whole genome assemblies (n = 183) and Illumina read sets (n = 419) from *L. rhamnosus* whole genome sequencing projects (Supplemental Table 1). Alignments of the four newly sequenced and 602 comparator genomes against the reference genome sequence of strain ATCC 11443 were used to perform a maximum likelihood (ML) phylogenetic analysis of the *L. rhamnosus* species and investigate the population context of the patient’s bloodstream isolate and the yogurt isolates (Fig. 2A).

**Figure 2.**
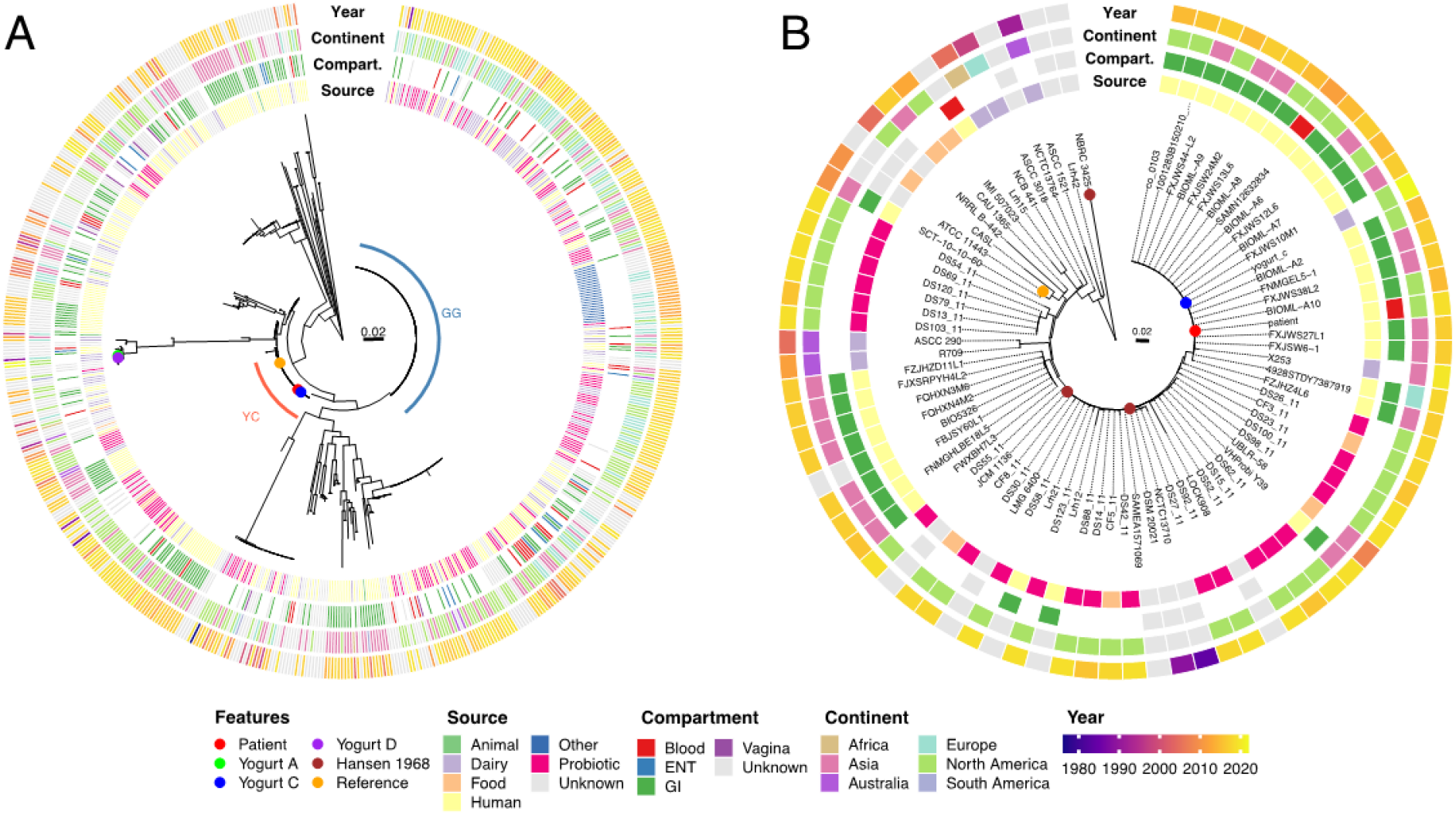
Whole genome phylogenetic analysis of *Lacticaseibacillus rhamnosus*. A) Maximum likelihood phylogenetic tree from whole genome alignment of sequences of the patient’s bloodstream isolate (“Blood” – red circle), representative yogurt isolates (“Yogurt A, C, or D” – green, blue, purple circles), and *L. rhamnosus* assemblies and sequence reads from the NCBI database (n = 611) to the reference genome sequence of strain ATCC 11443 (orange circle). Red arc indicates clade YC containing the patient and yogurt C isolates, blue arc indicates clade GG. B) Maximum likelihood phylogenetic tree of isolates sharing a clade with the patient’s bloodstream isolate and yogurt C (Clade YC, indicated by red arc in panel A). Brown circles indicate Hansen 1968 isolate sequences. In both panels, rings from inside to outside indicate, in order: 1. isolation source, 2. body site of isolation for human sourced isolates, 3. continent of isolation, and 4. year of isolation. Taxon labels in bold indicate sequences for which only genome assemblies were available. Scale bars indicate genetic distance.

In the phylogenetic analysis, the largest clade was comprised of 230 closely-related isolates represented by the GG strain of *L. rhamnosus* (Fig. 2A and Supp Fig. 1), a strain that is widely used in probiotic preparations [11]. Most GG sequences were isolated from dairy (n = 68, 30%) or probiotic product (n = 61, 26%) sources. Of the 59 clade GG isolates identified as originating from human samples, most were isolated from feces or other GI sources (n = 33, 14%) while 18 isolates (8%) were isolated from the blood (Supp. Fig. 1). Despite an abundance of both dairy and human bloodstream isolates in the GG clade in the public sequence database, neither the patient’s bloodstream isolate nor any of the three *L. rhamnosus* isolates from commercial yogurt samples A, C, or D were part of the GG clade. Two of the yogurt isolates, A and D, shared identical sequences with each other but were not closely related to any other sequences currently deposited in the NCBI database. In contrast, the patient’s isolate and the yogurt C isolate were both closely related to each other and were members of a separate clade, which we will refer to as clade YC, consisting of 80 isolate sequences (Fig. 2A, red arc).

Sequence alignment and ML phylogenetic analysis of the 79 isolates sharing a clade with the patient’s bloodstream and yogurt C isolate sequences revealed limited genomic diversity (Fig. 2B). Average pairwise genetic distance among the 81 isolates in clade YC was 32 single nucleotide variants (SNVs) (range 0 – 293), slightly higher than the average pairwise distance of 25 SNVs (range 0 – 261) among clade GG isolates and considerably lower than the average SNV count of 50,665 (range 38,566 – 51,649) between sequences across the GG and YC clades (Supp. Fig. 2). Unlike the GG clade, most of the isolates in clade YC were of human origin (n=34, 42%) with probiotic products as the next most common isolation source (n=22, 27%). Only seven isolates (9%), including yogurt C, were identified as originating specifically from dairy products. Within clade YC, just three isolates (4%), including the patient’s isolate, were identified as originating from the blood, whereas 31 isolates (38%) were isolated from human GI samples. None of the available metadata for the GI isolate sequences in the YC clade definitively indicated that any of the samples were obtained from patients in disease states (i.e. diarrhea), and most were documented as isolated from samples donated to stool banks (Supp. Table 1). Of note, several of the human GI isolates as well as the previously reported bloodstream isolate SAMN12632834 were identical (0 SNV differences) to the yogurt C isolates. The oldest sequence in this clade was isolated in 1984, but the majority of sequences were from isolates reported as collected between 2015 and 2019 (n=56, 69%). Isolates in clade YC were predominantly obtained in North America (n=38, 47%) and Asia (n=22, 27%) with three isolates from Australia and two from Europe. Of note, this clade also contained three sequences of the *L. rhamnosus* type strain “Hansen 1968” identified by different repository designations: DSM 20021, JCM 1136, and NBRC 3425 [12]. These three sequences were generated by different institutions and differed by between 1 and 220 total pairwise SNVs in their alignments. The two most closely related Hansen 1968 isolates, DSM 20021 and JCM 1136, differed from the yogurt C isolate by 10 and 11 SNVs, respectively. Together, these phylogenetic patterns suggest a diffuse geographic distribution of clade YC, possibly originating from the Hansen 1968 type strain, with very little or no genetic variation over time.

The bloodstream isolate differed from yogurt C isolate and several of the NCBI human GI tract isolates by only one SNV: an adenine to guanine amino acid change at position 1,753,277 relative to the sequence of the ATCC 11443 reference strain. This variation encodes a nonsynonymous amino acid substitution, M1A, disrupting the start codon of the NrdR transcriptional regulator gene (locus ID CEA83_RS08935). None of the 602 publicly available *L. rhamnosus* genome sequences nor any of the other yogurt isolate sequences differed from the reference sequence at this position.

## DISCUSSION

*Lacticaseibacillus* species are a rare cause of invasive bloodstream infections and is even more rarely associated with endocarditis [3–6]. Predisposing factors that could increase the risk of invasive infection with *Lacticaseibacillus* include immune compromising conditions or treatments and disruption of intestinal mucosal barriers, both of which are thought to allow these otherwise low pathogenicity organisms to establish infection. An open question remains as to whether consumption of *Lacticaseibacillus*-containing products such as yogurt or probiotics predispose an individual to invasive infection. In this study, whole genome sequencing confirmed that the patient’s bloodstream isolate belonged to the species *rhamnosus* and that this same species of *Lacticaseibacillus* was similarly found in three of four separate brands of commercially available yogurt purchased for this study. Further, whole-genome based analysis revealed that three randomly selected *L. rhamnosus* isolates cultured from one of the yogurt brands (yogurt C) were nearly identical to the patient’s bloodstream isolate, differing only at a single nucleotide position over more than 2.9 million aligned bases. However, this study also revealed several sequences in the public database were genetically identical to the yogurt C isolate and equally closely-related to the patient’s bloodstream isolate. The lack of genetic diversity between these strains precludes identification of the source of this patient’s bloodstream infection, but two possibilities exist. First, invasive infection could have resulted directly from recent yogurt consumption by the patient leading to gut translocation of strains from the consumed product. Alternatively, it is possible that the patient’s gut may have been previously colonized by the strain in question from earlier ingestion and infection arose from translocation of the patient’s own GI flora. It is noteworthy that the sequences most closely related to the patient’s isolate were largely of human GI origin, were identified from individuals around the world, and most were not identified as having been cultured from dairy or probiotic products. Additionally, we describe here a *L. rhamnosus* strain cultured from yogurt that is genetically identical to these strains. This suggests that this strain of *L. rhamnosus* of yogurt origin is capable of colonizing the human GI tract, though whether this is transient or long-term colonization cannot be determined from the available data. In addition to their identical or nearly identical genetic sequences, the geographic and temporal diversity of clade YC isolates would suggest that they stem from a common origin or source before being globally distributed. It appears likely that one or more commercial dairy or probiotic products may be produced using this particular strain of *L. rhamnosus* and that, subsequently, the identical strain can be recovered from the GI tracts of individuals who consume such products [13–22].

Similarly, the available public sequence records and associated metadata suggest that GG clade isolates of *L. rhamnosus* have been recovered from the human GI tract as well as bloodstream infections. *L. rhamnosus* GG (LGG) was first isolated from fecal samples of a healthy adult and has since been widely used as a probiotic strain largely due its acid and bile resistance, favorable growth profile, and intestinal epithelial adhesion properties [13]. LGG has been associated with invasive disease as evidenced by published case reports and bloodstream isolate sequence data [14–18]. Review of the sequence database also revealed that, like the clade YC of *L. rhamnosus* described here, GI carriage of LGG clade isolates can occur. This is supported by prior studies demonstrating recovery of *Lacticaseibacillus* GG in the GI tract or stool during or after probiotic therapy, though GI carriage was frequently not durable for more than a few weeks after ceasing ingestion of the *Lacticaseibacillus*-containing probiotic [19–22]. Notably, study sizes in these reports were small, but suggest that long-term colonization by dairy or probiotic strains of *L. rhamnosus* may depend on repeated consumption of strain-containing products. Finally, the isolation of genetically identical *L. rhamnosus* isolates from two readily obtained yet different brands of store-bought yogurt (A and D) that were distinct from any other sequences in the public database, suggests that non-LGG commercial strains of *L. rhamnosus* are underrepresented in public databases. As WGS of bacteria isolated from humans becomes more cost-effective and widespread, epidemiologic and clinical conclusions will increasingly be drawn from comparisons of such isolates to sequences present in public databases. Therefore, it would be of great benefit for a more complete picture of potentially invasive *L. rhamnosus* strains used in commercial products to be defined. This study adds to public knowledge of genetic sequences of *L. rhamnosus* isolates consumed by humans that also have pathogenic potential.

We identified only a single nucleotide difference in the patient’s bloodstream isolate relative to the yogurt C isolate and other previously reported GI isolate sequences in the YC clade. This nucleotide change resulted in a nonsynonymous amino acid change disrupting the start codon of the NrdR transcriptional regulator gene. The *ndrR* gene is a transcriptional repressor usually found clustered with ribonuclease reductase (RNR) genes or genes involved in primosome assembly and DNA replication in many bacterial species [23–25]. NdrR is thought to interact with thioredoxin and play a role in maintenance of homeostasis in bacteria [26]. Despite this role, prior studies in other species of bacteria have shown that deletion of *ndrR* did not result in alteration of growth profiles in culture, though deletion of this gene in *Pseudomonas aeruginosa* and *Streptococcus pyogenes* did result in variability in adherence of the bacteria to host cells [27, 28]. It is unclear whether this mutation in the patient isolate provided any selective advantage in prolonged bloodstream infection or endocarditis or merely represents neutral evolutionary variation.

Our analysis suffers from several limitations. To try to capture the breadth of diversity in the *L. rhamnosus* population structure we sought to include as many high-quality genomic sequences as possible. By including whole-genome sequence assemblies, the variety of methods used by different groups to generate whole genome assemblies could have introduced and/or compounded errors in the deposited data, possibly artifactually changing the measured genetic distance and relatedness of these isolates. We sought to minimize the effect of such errors on our results by using a pseudoread approach to employ a single shared alignment and SNV-calling method for both sequence reads and assembly data sets. We have highlighted the different input sequence types in Figures 2B and Supp. Fig 1. We did not have concurrent GI samples collected from this patient to examine possible GI carriage of the same strain and we also were not able to determine definitively the specific brands of yogurt the patient preferred. Intriguingly, one of the brands of yogurt the patient consumed as remembered by family members was included in this study (yogurt A). Isolates from yogurt A did not match the bloodstream isolate sequence, though the family’s confidence in recalling the patient’s preference was not high. Likewise, detailed information on the diversity of *L. rhamnosus* strains included in probiotic and other consumable products is lacking. Without such information, the true incidence of sustained GI carriage of commercial strains in the human population is unclear. Therefore, the risk of invasive infection from prior commercial strain colonization remains difficult to determine.

## Materials and Methods

### Bacterial strains and growth conditions

The patient’s isolate was obtained from a blood culture drawn during the patient’s second hospitalization for recurrent *Lactobacillus* bacteremia. Single servings of commercial yogurt brands were purchased from a local store of a national supermarket chain and stored at 4° C. Samples were streaked onto De Man, Rogosa and Sharpe (MRS) *Lactobacillus-*selective agar (Research Products International, IL) and grown for a minimum of 48 hours at 37° C in anaerobic conditions.

### Whole-genome sequencing

Isolates were grown overnight in MRS broth topped with paraffin oil without shaking at 37°C to create microaerophilic growth conditions. DNA extraction was performed using a QIAamp BiOstic Bacteremia DNA Kit (Qiagen) according to the manufacturer’s instructions. Sequencing libraries were prepared with a plexWell™ 96 kit (seqWell) and sequenced on an Illumina MiSeq platform (Illumina, Inc., San Diego, CA) using a v3 reagent kit yielding 2 x 301 bp paired-end reads totaling 6.4 Gbp of sequence, with an approximate per-sample read coverage of 92-fold. Adapter sequences were removed and reads were quality trimmed using fastp v 0.32.2 [29] and then *de novo* assembly was performed with SPAdes v3.15.4 [30]. Assemblies were filtered to remove contigs smaller than 200 bp or with average read coverage of less than 5. Sequences were deposited in the National Center for Biotechnology Information (NCBI) database (BioProject accession number PRJNA908878).

### Public sequences

To identify available *L. rhamnosus* genome assemblies from the NCBI database, all genome records with genome ID 913 as of April 20, 2022 were identified from the NCBI Genome database and nucleotide sequences were downloaded from the GenBank FTP server. Of the 246 *L. rhamnosus* genome sequences identified, 15 were excluded as low quality for assembly contig counts > 400 and/or exceeding two standard deviations (220,627 nt) above or below the average total contig count of the entire sequence set (2,968,492 nt). To identify *L. rhamnosus* sequencing read sets, the NCBI Sequence Read Archive (SRA) database was searched for all records fulfilling the following Entrez query: “Lacticaseibacillus rhamnosus”[organism] AND “WGS”[strategy] AND “Illumina”[platform]. As of May 9, 2022, this query identified 495 records in the SRA database. Of these records, 49 sets were excluded for consisting only of single-ended reads. Read sets were further filtered to randomly select one set of reads from multiples sharing a single BioSample ID (n = 15). Two read sets with read name errors in the downloaded files were also excluded. Eight read sets were identified by 16S rRNA gene sequence to be from species other than *L. rhamnosus* and were excluded. Finally, two read sets with genome alignment coverages of less than 80% of the reference genome length were removed to leave a total of 419 paired read sets for analysis. Accounting for overlaps between the assembly and read sets in which both an assembled genome and read sets were deposited under the same BioSample record (n=39) or assembly and sequence reads for the same isolate were deposited under two separate BioSample records (n=9), the total number of sequences of unique *L. rhamnosus* isolates obtained from the NCBI GenBank and SRA databases was 602 (Supplemental Table 1). Metadata for each assembly and read record, including host, isolation source, collection date, and geographic location of isolation, were obtained from the NCBI BioSample database when available.

### Single nucleotide variant identification

For sequences with only genome assemblies available, the assembly sequences were used to generate pseudoread sets for alignment. Sequence reads and assembly pseudoread sets were aligned to the reference genome sequence of ATCC 11443 (NCBI accession GCA_003433395.1) using bwa v 0.7.15 [31]. Strain ATCC 11443 was chosen as an alignment reference as it is a complete ungapped genome assembly and closely related to the patient isolate. Single nucleotide variants relative to the reference were identified using bcftools v1.9 skipping bases with base quality lower than 25, alignment quality less than 30, and using a haploid model. Variants were further filtered as previously described [32] using the bcftools_filter software (https://github.com/egonozer/bcftools_filter) to remove variants with single nucleotide variant (SNV) quality scores less than 200, read consensuses less than 75%, read depths less than five, read numbers in each direction less than one, or locations within repetitive regions (as defined by blast alignment of the reference genome sequence against itself).

### Phylogenetic analysis

For the total set of all consensus sequences, variant positions with base calls in less than 100% of isolate sequences (i.e. non-core positions) were masked with N’s using ksnp_matrix_filter.pl [33]. Separate complete sequence alignments consisting only of consensus sequences from isolates in the same clade as the patient’s isolate (YC) or those in clade GG (Supplemental Table 1) were generated. Maximum likelihood phylogenetic trees were created from genome alignments with IQ-TREE v1.6.1 [34, 35]. Tree topology was assessed with the Shimodaira-Hsegawa approximate likelihood ratio test (SH-aLRT) [36]. Phylogenetic tree visualization and annotation was performed using R and the ggtree package [37, 38].

## ACKNOWLEDGEMENTS

The work was supported by the National Institute of Allergy and Infectious Diseases at the National Institutes of Health U19AI135964 [E.A.O and A.R.H] and R01AI118257, K24AI104831, R21AI153953, and R21AI164254 [A.R.H.] and an American Cancer Society Clinician Scientist Development Grant #134251-CSDG-20-053-01-MPC [K.E.R.B]. This work was supported by the Northwestern University NUSeq Core Facility which is supported, in part, by the NCI CCSG P30 CA060553 award to the Robert H. Lurie Comprehensive Cancer Center. This work was also supported in part through the computational resources and staff contributions provided by the Genomics Compute Cluster, which is jointly supported by the Feinberg School of Medicine, the Center for Genetic Medicine, and Feinberg’s Department of Biochemistry and Molecular Genetics, the Office of the Provost, the Office for Research, and Northwestern Information Technology. The Genomics Compute Cluster is part of Quest, Northwestern University’s high-performance computing facility, with the purpose to advance research in genomics. The funders had no role in study design, data collection and analysis, decision to publish, or preparation of the manuscript. We would like to thank members of the Center for Structural Genomics of Infectious Diseases (CSGID) and the Hauser laboratory for their valuable comments during numerous discussions of this work.

---

**Supplemental Figure 1.**
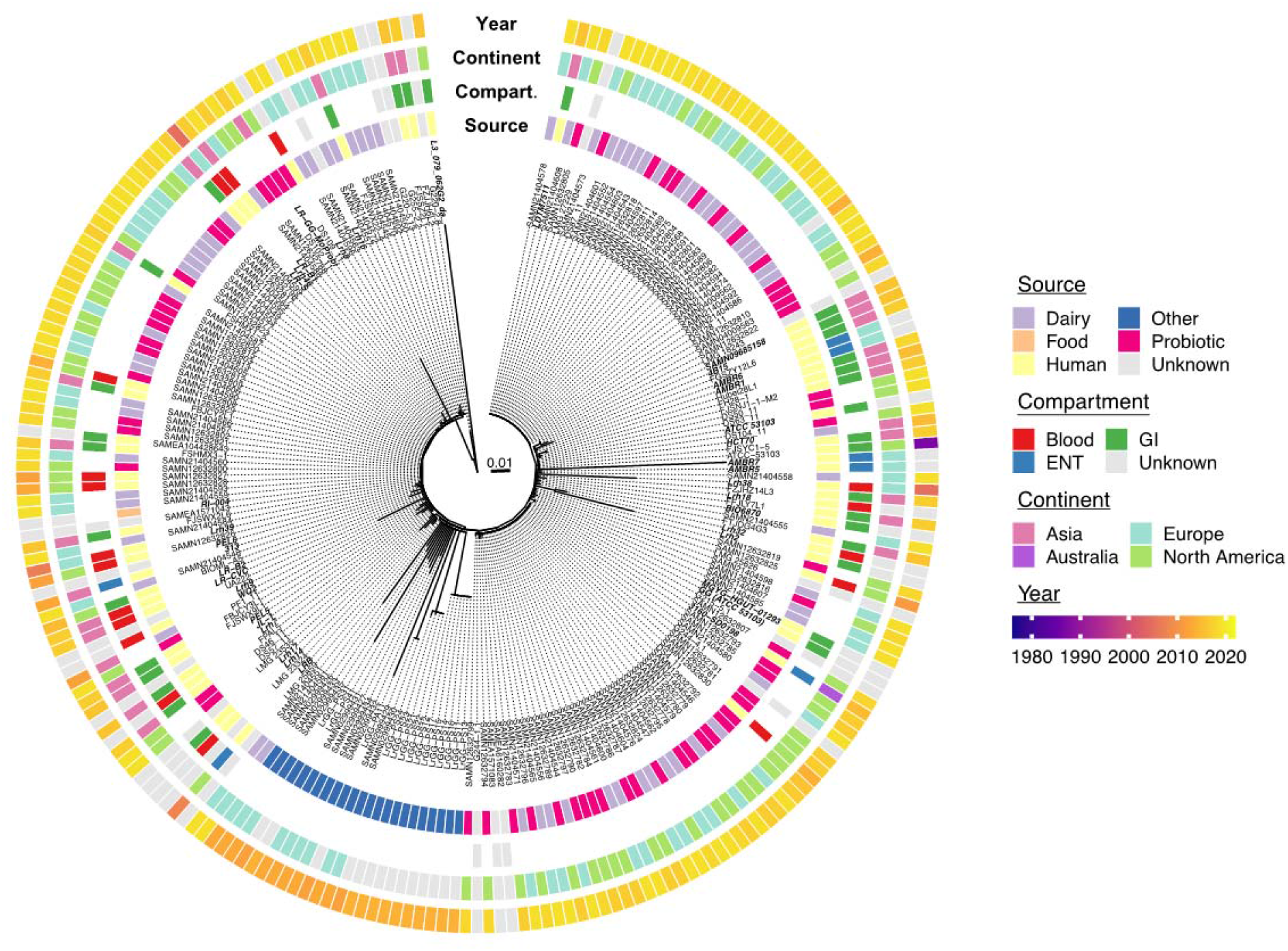
Maximum likelihood phylogenetic tree of clade GG isolates. Maximum likelihood phylogenetic tree from whole genome alignment of clade GG isolates (indicated by blue arc in Figure 2A) to the reference genome sequence of strain ATCC 11443. Rings from inside to outside indicate, in order: 1. isolation source, 2. body site of isolation for human sourced isolates, 3. continent of isolation, and 4. year of isolation. Taxon labels in bold indicate sequences for which only genome assemblies were available. Scale bar indicates genetic distance.

**Supplemental Figure 2.**
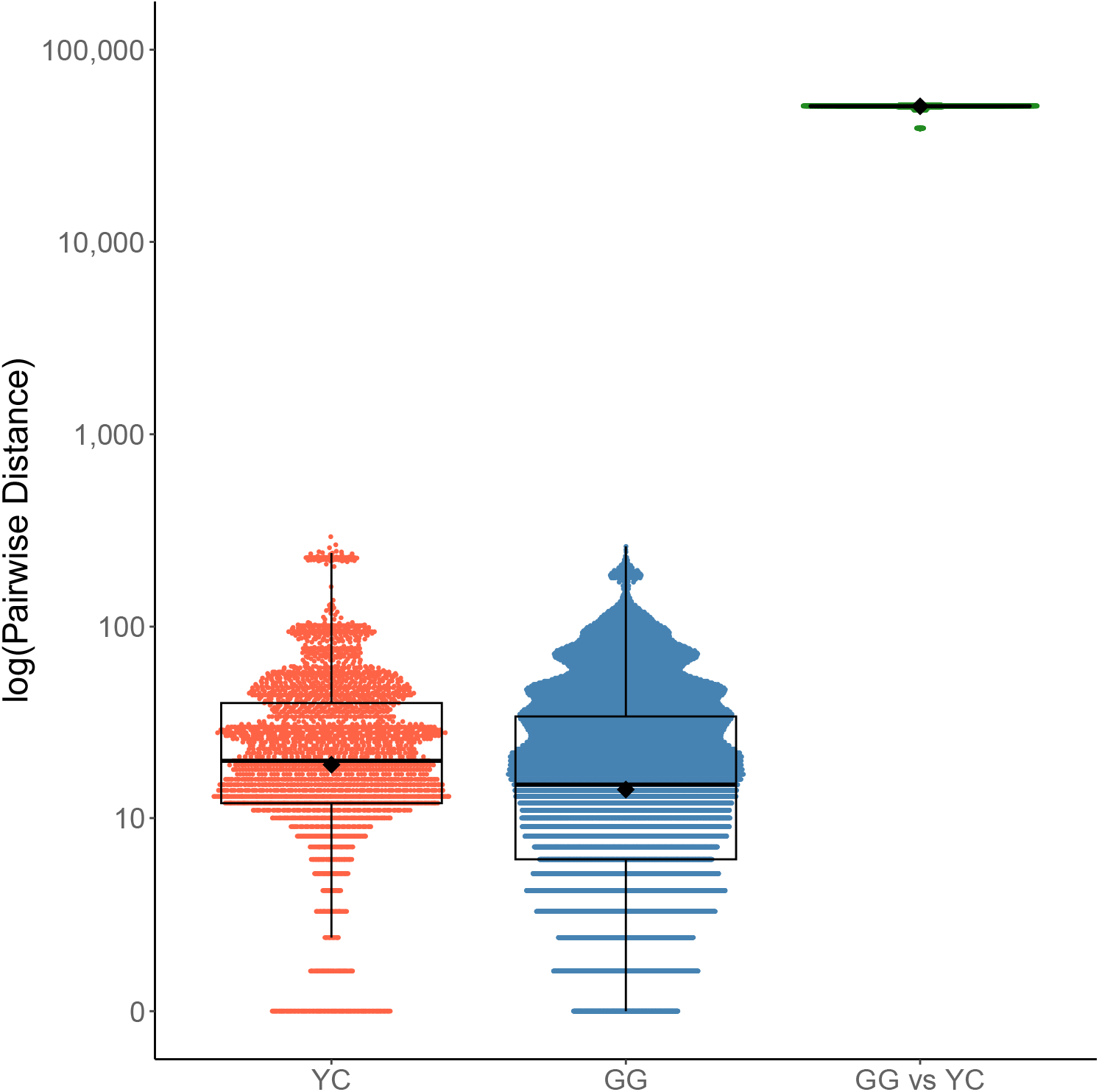
Pairwise genetic distances between isolates in the GG and YC clades. Dots indicate all pairwise genetic distances as measured by single nucleotide variant (SNV) differences among the 81 sequences in the clade including the patient’s bloodstream isolate and the yogurt C isolate (“YC”, red), among the 230 clade GG isolates (“GG”, blue) or between the two clades (“GG vs YC”, green). Overlying box and whisker plots indicate median SNV counts (thick black line) as well as first and third quartiles. Whiskers indicate 1.5 x the interquartile range (IQR) and average SNV counts per group are indicated by black diamonds. Y-axis is plotted on the log_10_ scale.

